# The Neural Basis of Event Segmentation: Stable Features in the Environment are Reflected by Neural States

**DOI:** 10.1101/2024.01.26.577369

**Authors:** Djamari Oetringer, Dora Gözükara, Umut Güçlü, Linda Geerligs

## Abstract

Our senses receive a continuous stream of complex information. Parsing this information into meaningful events allows us to extract relevant information, remember it, and act upon it. Previous research has related these events to so-called ‘neural states’: temporally and regionally specific stable patterns of brain activity, which tend to coincide with events in the stimulus. Neural states show a temporal cortical hierarchy: short states are present in early sensory areas, while longer states can be found in higher-level areas. Here we investigated what these neural states represent. We hypothesized that states at different levels of the cortical hierarchy are shaped by aspects of the stimulus to which these brain areas are responsive. To test this hypothesis, we analyzed fMRI data of participants watching a movie, using a data-driven method to identify the neural states. We found support for the aforementioned hypothesis: specifically the parahippocampal place area and retrosplenial cortex, known to be sensitive to places, showed an alignment between neural state boundaries and moments in the movie with a change in location, independent of changes in visual features and other covariates. These findings suggest that neural states reflect stable features in the (internal model of) the external environment, and that the cortical temporal hierarchy partly reflects the temporal scales at which representations of the environment evolve.

## 1 Introduction

The continuous stream of sensory input that we receive in our daily lives contains a large amount of information. Despite this complexity, we are seemingly easily able to process this information and use it appropriately. This phenomenon can partially be explained by the cognitive skill of event segmentation, which entails that we automatically segment incoming information into distinct events (Newtson et al., 1977; Zacks, 2020; Zacks et al., 2007). Here, events are “segments of time at a given location that is conceived by an observer to have a beginning and an end” (Zacks & Tversky, 2001, p. 3). Event segmentation can support the processing of complex information by creating smaller chunks of meaningful information (Kurby & Zacks, 2008). It can additionally aid memory encoding by updating representations of the current event at event boundaries, freeing information from working memory (Kurby & Zacks, 2008; Zacks et al., 2007), and by increasing access to relevant information and reducing interference (Shin & DuBrow, 2020). Various behavioral studies suggest that events can be segmented at different levels of granularity and that coarse-grained event boundaries tend to co-occur with fine-grained boundaries (Kurby & Zacks, 2008; Newtson, 1973; Zacks, Tversky, & Iyer, 2001). These results suggest that events are structured as a nested hierarchy, where big events contain multiple smaller ones.

While the behavioral properties and consequences of event segmentation have been studied extensively, much less is known about the neural mechanisms underlying event segmentation. Initially, the neural underpinnings were studied by looking at evoked responses at the moment of an event boundary, while participants experienced a naturalistic and dynamic stimulus. Event boundaries have been associated with an increased neural response in a wide network of brain regions, including the precuneus, fusiform gyrus, anterior and posterior cingulate gyrus, temporal gyrus, and middle frontal gyrus (Speer et al., 2007; Zacks, Braver, et al., 2001; Zacks et al., 2010). A number of studies have particularly looked into the event boundary response in the hippocampus, which is larger after events that are later remembered compared to those that are later forgotten (Ben-Yakov & Dudai, 2011; Ben-Yakov et al., 2013; Ben-Yakov & Henson, 2018), which further supports the importance of event segmentation for memory. These studies show where in the brain the neural activity is associated with the occurrence of event boundaries but they do not reveal how segmentation might be implemented at different levels of the cortical hierarchy. This gap can be addressed by studying so-called neural states, which are relatively stable patterns of brain activity within a specific region (Geerligs et al., 2021). Such stable patterns have previously been referred to as events in the brain (Baldassano et al., 2017), brain states (van der Meer et al., 2020), or neural events (Sava-Segal et al., 2023). To clearly distinguish information from neural activity and information from the stimulus and behavior, here we will be using neural states or neural state boundaries to refer to the stable patterns in the brain, and events or event boundaries to refer to units of information in the external environment, which are defined independently from brain activity data. Although events and neural state boundaries have been shown to overlap in time, they remain distinct concepts that should be considered separately.

One important characteristic of neural states is that they can be identified in local areas all throughout the cortex, making it possible to relate them to cortical hierarchies. Different approaches have also demonstrated temporal hierarchies across the brain. Using fMRI and a movie stimulus, Hasson et al. (2008) introduced the concept of temporal receptive windows (TRWs): the duration of the period in the past in which information affects the current response. This time window was longer in higher-level areas than early visual areas, showing a temporal hierarchy across the cortical hierarchy. These results have been replicated in the auditory domain (Lerner et al., 2011), and in electrophysiological measurements (Honey et al., 2012). More recently, Hasson et al. (2015) proposed the Hierarchy of Process Memory Framework, which emphasizes the ability of all cortical circuits to accumulate information over time, with the time duration depending on the temporal receptive window.

In their groundbreaking study on neural mechanisms of event segmentation, Baldassano et al. (2017) builds upon this Hierarchy of Process Memory Framework and relates it to neural states by proposing that each brain area along the cortical hierarchy segments information, starting with short neural states in low-level sensory areas and long states in high-level multimodal areas. In support of this theory it has now been shown that neural states form a partially nested hierarchy across a set of regions (Baldassano et al., 2017; Geerligs et al., 2022), similar to how fine-and coarse-grained events are structured in a nested hierarchy. Neural state boundaries have been shown to coincide with event boundaries, suggesting that neural states are driven by changes in the mental model or the external environment of the participant (Baldassano et al., 2017; Geerligs et al., 2022). Indeed, neural state durations can vary substantially over time within a particular brain region, suggesting that the durations do not reflect a fixed temporal scale of information processing but instead change dynamically based on temporal fluctuations in the features of the stimulus (Geerligs et al., 2022).

Even though it is clear that, in some brain regions, neural states are associated with events, it remains unclear what neural states in different brain areas actually represent. In this study, we tested the hypothesis that neural state boundaries in a given brain region reflect a change in the environment that is relevant for that brain region. Although this idea has been proposed before by Baldassano et al. (2017), and it is in line with the observed overlap between neural state and event boundaries, it has never been tested directly for other environmental features. As a conceptual example, the fusiform face area (FFA) would have a neural state boundary when you move your head to look from one person to another. If this is the case, neural states might represent stable features in the external environment or in the mental representation that a person has about that environment. Since aspects that are relevant for higher-level regions also tend to evolve at a slower timescale, this could also explain the observed temporal hierarchy in neural state durations. It should be noted that this hypothesis is not in line with the idea of an event boundary as an all-or-none global update of event representations (Zacks et al., 2007), but would rather suggest that there are incremental updates to situation models in different brain regions.

An alternative hypothesis is that the duration hierarchy of neural states could mostly be shaped by the intrinsic temporal receptive windows of each area across the cortex. In line with this possibility, intrinsic timescale and power spectra gradients have been observed even in the absence of a stimulus (Honey et al., 2012; Raut et al., 2020; Stephens et al., 2013). In this case, the previously observed alignment between event boundaries and neural states would not translate to other stimulus features. Such a unique nature of event boundaries in resetting the brain signal would instead be in line with global updating models of event segmentation (Zacks et al., 2007).

To investigate our hypothesis, we used an open fMRI dataset in which subjects were watching a movie. We investigated the neural states in a small set of regions of interest (ROIs) for which we could make clear predictions about the features in the external environment that they are responsive to. These were the parahippocampal place area (PPA) and retrosplenial cortex (RSC) that are both known to play an important role in visual scene recognition (R. A. Epstein & Higgins, 2007; R. Epstein & Kanwisher, 1998; Kriegeskorte & Kievit, 2013). If neural state boundaries in a particular region reflect a change in the environment that is relevant for that region, we expect that the RSC and PPA would show an alignment between their neural state boundaries and moments in the movie at which there was a change in location. Importantly, we expected this alignment between neural states and locations to be specific to these location-sensitive areas, and to be independent from co-varying changes in the stimulus such as event boundaries, and changes in low-level visual and auditory features. Therefore, we included a correction for covariates, and additionally investigated the neural state boundaries in an early visual ROI, where we expected to see an alignment between state boundaries and changes in low-level visual features, but not with location changes. The specificity of these associations was further investigated in a set of exploratory whole-brain analyses. We indeed observed a reliable and regionally specific alignment between neural state boundaries and changes in the stimulus that are associated with the region, independent of a multitude of covariates in the stimulus. These results indicate that neural states reflect stable features in the (internal model of) the external environment. They also suggest that the cortical temporal hierarchy in terms of neural state durations partly reflects the temporal scales at which different aspects of the environment naturally evolve.

## 2 Methods

### fMRI data

To investigate neural states during complex and dynamic input with some available stimulus annotations, we used an open dataset called StudyForrest (www.studyforrest.org/). We specifically used the fMRI dataset in which 15 participants (age 19–30, mean 22.4, 10 female, 5 male) watched a slightly cut-down version of the movie Forrest Gump dubbed in German (Hanke et al., 2016). The 2-hour movie was divided into 8 segments that were presented across two sessions on the same day. In total there were 3599 volumes of data acquired with a 32-channel head coil using a whole-body 3 Tesla Philips Achieva dStream MRI scanner. The acquired images were T2*-weighted echo-planar images (gradient-echo, 2 s repetition time (TR), 30 ms echo time, 90^◦^flip angle, 1943 Hz/Px bandwidth, parallel acquisition with sensitivity encoding (SENSE) reduction factor 2). There were 35 axial slices (thickness 3.0 mm) with 80 ×80 voxels (3.0 × 3.0 mm) of in-plane resolution, 240 mm field-of-view (FoV), and an anterior-to-posterior phase encoding direction with a 10% inter-slice gap, recorded in ascending order. For anatomical alignment, we used the T1-weighted high-resolution structural scans as provided by Hanke et al. (2014).

To functionally define our regions of interests, we used the functional localizer data that is additionally part of StudyForrest (Sengupta et al., 2016). The MRI acquisition setup of this localizer was the same as for the movie data.

### 2.1 Annotations

In order to investigate the relationship between neural states and the stimulus, a multitude of stimulus annotation categories were gathered: shots, locations, events, auditory information in the form of Mel-Frequency Cepstral Coefficients (MFCC), speech, low-level visual features, and visual features of different levels of complexity using a convolutional neural network. Annotations of shots and locations were taken from Häusler and Hanke (2016), who manually annotated the data by determining when there was a new shot on a frame-by-frame basis, and coding at which location each shot took place. They defined three levels of locations: ‘locale’ (e.g. ‘living room’ or ‘corridor downstairs’), ‘setting’ (e.g. ‘Gump house’ or ‘Jenny’s house’), and ‘major location’ (e.g. ‘Greenbow Alabama’ or ’California’). In our analysis, we only used ‘locale’ (from this point on referred to as ‘small-scale locations’) and ‘setting’ (from this point on referred to as ‘large-scale locations’). The level ‘major location’ was not included as it reflects conceptual changes that are probably more relevant to other areas, such as the anterior temporal lobe (Bonner & Price, 2013). Further, this influence, if present, would depend on the subjects’ knowledge of the geography of the United States of America, while the subjects were German. Finally, given the scale of this annotation, the number of changes in such major locations was relatively low throughout the movie, making it more difficult to detect any significant effect, as is often an issue in naturalistic stimuli with variables that have a naturally low rate of occurrence (Hamilton & Huth, 2020).

To ensure that associations between neural state boundaries and location changes were not caused by changes in other aspects of the stimulus, we also acquired annotations of low-level visual features, MFCC, speech, and event boundaries that could be included in the analysis as covariates. The annotations of low-level visual features were based on perceptual hashes (using *ImageHash* version 4.1.0) of the various frames and were taken from the StudyForrest GitHub pages. Perceptual hashing is an algorithm that takes an image as input, reduces size and color, extracts low-frequency information using a Discrete Cosine Transform, and outputs a string of characters that can be interpreted as the fingerprint of that image. This algorithm is designed such that the fingerprints of images are close to each other when they contain similar features, and is therefore insensitive to cropping or a change in resolution (see e.g., Fridrich & Goljan, 2000; Zauner, 2010). Specifically relevant in a movie, this measure is insensitive to ongoing movement. For example, two consecutive frames in which a character is running towards the left would be seen as very similar to the human eye, as well as to perceptual hashing, while a pixel-based measurement would be very sensitive to such movement. Instead, this algorithm is sensitive to the overall layout and shapes, and thus to the gist representation of an image rather than local edges and high-frequency details. The difference in low-level visual features over time were defined on a frame-by-frame basis: after computing each frame’s perceptual hash, the Hamming distances between the perceptual hash values of consecutive frames were measured, and consequently normalized.

The perceptual hashes described above only account for the visual features that are of one specific level of complexity, while visual features of other levels of complexity also have the potential to drive associations between for example changes in location and neural state boundaries. Therefore, visual features at various levels of complexity were additionally extracted from the movie by using Alexnet (Krizhevsky et al., 2012): a convolutional neural network trained for object recognition. Each stimulus frame, excluding the gray bars at the bottom and top of the screen, was given as input to Alexnet with pretrained weights. The outputs of all layers except the output layer were extracted, giving us an approximation of visual features across 8 layers, which can be seen as 8 different levels of complexity (Zeiler & Fergus, 2014). As we were interested in the frame as a whole, we averaged the outputs over the spatial dimensions to discard the spatial information. Then, the frame-to-frame differences were computed by taking the cosine distance between the resulting vectors of two consecutive frames, and these differences were scaled to have a maximum of 1 per layer.

To include auditory information in the analyses, we extracted MFCCs from the original stimulus per run. The coefficients were first averaged per fMRI volume (after having added a delay; see below), and then converted to changes in MFCC by computing the cosine distance between consecutive volumes. These distances were then scaled such that the maximum value was set to 1. We additionally defined the speech onsets and offsets based on word onset and duration information from previously defined annotations (Häusler & Hanke, 2021). The start of each silence period and each speech period (except for the very first timepoint of the run) was considered a ‘change in speech’, giving us a timeline of ones and zeros. Here, ‘silence’ was defined as a period of no words having been spoken for one TR (i.e., two seconds). Thus, each 1 in this timeline represents the onset or offset of speech.

Finally, event boundaries of the movie stimulus were defined by Ben-Yakov and Henson (2018), whose subjects watched the Forrest Gump movie stimulus and indicated with a button press when they thought a new event had started. These subjects were native English and watched the movie in either German or English, though the button presses were not significantly different between the original English movie and the German dubbed version. 0.9 seconds were subtracted from each button press to account for response time, and then button presses of different subjects were taken together when they were close in time. In order to exclude accidental presses and event boundaries with low between-subject reliability, only moments at which at least 5 out of 16 subjects had pressed a button around roughly the same time were considered an event boundary, giving us a binary timeline of an event boundary being present at a certain timepoint or not.

Overall, the correlations between the different annotations were high (see Figure S2). This is due to the nature of movies and the annotation categories. For example, a change in location can only occur when there is also a new shot. Similarly, a change in large-scale location (e.g. from a house to a school) always coincides with a change in small-scale location (e.g. from the living room to the classroom). Therefore, the small- and large-scale locations have a nested relationship. Although the event boundaries were acquired with a more subjective method, the moments at which observers press the button generally tends to coincide with changes in location (Magliano et al., 2001). For more descriptives and examples of the annotations, see Supplementary section A.

### 2.2 Preprocessing

In this study we used the movie fMRI data that had previously been partially preprocessed and denoised by Liu et al. (2019). They performed motion correction, slice timing correction, brain extraction and high-pass temporal filtering (cutoff of 200 s). They then denoised the data by using spatial ICA decomposition and manual inspection of the resulting components to remove artifacts as a result of hardware, head motion, physiology, and unclear sources. As the computation of neural state boundaries requires spatial information, we specifically used the unsmoothed version of this published dataset.

We further preprocessed the data using SPM12 software (www.fil.ion.ucl.ac.uk/spm) by aligning the functional data between the eight runs, performing coregistration with the T1 anatomical image of each subject, followed by segmentation, and finally normalization (Ashburner & Friston, 2005) to MNI space. Only gray matter voxels were included, which were defined as voxels that survive the threshold of 0.35 in the gray matter probability map (TPM) as provided by the SPM software. The final gray matter mask only contained voxels that were measured for all subjects in all runs. Here, a voxel was labeled as ‘measured’ during one run when its value during the first timepoint of that run was higher than 0.8 times the mean of all voxels at that same timepoint.

After normalization and gray matter masking, the data was z-scored per voxel per run. To functionally align the subjects’ data and thereby improving inter-subject correlation, hyperalignment was performed on the whole brain, using the implementation of the PyMVPA toolbox (Guntupalli et al., 2016; Haxby et al., 2020). This hyperalignment was computed on the fMRI data of run 4 (488 volumes) using searchlight-hyperalignment. The searchlights had a radius of 3 voxels and all voxels within the gray matter mask were used as the center of a searchlight. The computed hyperalignment parameters were then applied to all remaining runs. Since run 4 was used to compute the transformation parameters, as well as the optimal delay between annotations and neural states (see below), this run was excluded from any further analysis. Run 4 was chosen as it contained a relatively low number of changes in location in the movie, and because it was roughly in the middle of the movie, making it more representative than for example the first or last run.

### 2.3 ROI definitions

To functionally define the ROIs, we used data from a functional localizer experiment that was part of the StudyForrest dataset (Sengupta et al., 2016). In this experiment, the StudyForrest participants that had also participated in the audio-visual movie part of the dataset were presented with faces, objects, houses, landscapes, bodies and scrambled images while fMRI data was collected. After data preprocessing, Sengupta et al. (2016) performed a GLM analysis with one regressor per image category. Next, they computed multiple contrasts per subject to find subject-specific voxels that are sensitive to one or multiple of the aforementioned image categories. We used some of these contrasts to define our ROIs. For the left and right PPA and the RSC, we specifically used the contrast of houses and landscapes with all other image categories. For left and right early visual ROIs we took the contrast of scrambled images with all other image categories, and we excluded the midline, defined as a sagittal plane with a thickness of 2 voxels surrounding the exact middle of the left-right axis, to be able to separate a left and right cluster. To obtain group-level ROIs, required for obtaining reliable neural states, we took the subject-level contrast values, normalized them to MNI space using SPM12, and applied a between-subject t-test per voxel. We then applied the gray matter mask and thresholded the resulting t-values such that one cluster of about 100 voxels was left at roughly the expected anatomical location (Nasr et al., 2011). Descriptives and visualizations of the resulting ROIs can be found in Supplementary Section B.

### 2.4 Analysis

To reliably identify neural states using Greedy State Boundary Search (GSBS; Geerligs et al. 2021), the data must have a high enough signal-to-noise ratio (SNR). Since the stimulus is only presented once, data from a single subject has low SNR, which is why analyses were performed on the group-level data by averaging the data across participants. To obtain the neural states per ROI, we applied GSBS to the group-averaged ROI data in each run. GSBS is a data-driven method to extract the neural states from fMRI data in an iterative manner, by trying out a new boundary at each possible timepoint and selecting the one that gives the best fit. The algorithm determines the optimal number of states by maximizing within-state correlations and minimizing between-state correlations. Parameters were set to the values as recommended by Geerligs et al. (2021): the maximum number of neural states was set to half the number of time points in a given run, and finetuning was set to 1, meaning that each neural state boundary could be shifted by 1 TR after each iteration, in case that would give a better fit. We specifically used the states-GSBS algorithm, which is an improved and more reliable version of GSBS, introduced by Geerligs et al. (2022), that potentially places two boundaries in a given iteration rather than always placing one boundary at a time. Taken together, this is quite different from the method introduced by Baldassano et al. (2017), who used a Hidden Markov Model (HMM). Although both are valid methods to identify neural state boundaries, GSBS is more appropriate here as the HMM has lower performance when the neural states are of variable durations (Geerligs et al., 2021), which is applicable in this study as we investigate the relationship with stimulus features that can vary in duration throughout the movie (such as locations). Additionally, GSBS gives us the benefit of identifying the number of neural states in a data-driven approach, without having to re-run the algorithm many times.

GSBS gave us a timeline of neural state boundaries per run per ROI, including the strength of each boundary, with zeros indicating the absence of a neural state boundary and any value above zero indicating the presence as well as the strength of a boundary. The strength of a boundary is defined as 1 minus the Pearson correlation of the average neural activity patterns of the two neural states that surround the boundary. Thus, a higher strength implies that the two consecutive neural states are more dissimilar. To compare the neural state boundaries to the various annotations, all annotations were first delayed by 4.5 seconds to account for the hemodynamic response, and then downsampled from seconds to fMRI volumes. The delay of 4.5 seconds was based on an independent analysis with the data from the unused run 4, as shown in Supplementary Section C. Finally, all values per run were concatenated, creating one vector per annotation for the whole movie, as well as one neural state boundary vector per ROI.

To measure the overlap between the neural state boundaries and various annotations, we used partial Pearson correlation. Given the high correlations between the various annotation categories, partial correlation was necessary to minimize the possibility that a significant correlation could be explained by a covariate instead of the annotation of interest. For example, a significant correlation with changes in location could actually be due to changes in low-level visual features or vice versa. By using partial correlation, any association we observe cannot be explained by any of the covariates used in the analysis. Partial correlation was computed between the neural state timeline of a particular brain area and the timeline of an annotation of interest, while correcting for appropriate covariates (see below). In the neural state timeline, each zero represented the absence of a boundary, and each value above zero indicated the presence of a neural state boundary including its strength with a maximum of 2. The timeline of the annotation consisted of either binary or continuous values depending on the annotation of interest. It should be noted that the resulting correlation coefficients may not be directly comparable between brain areas to measure a difference in alignment, as extra neural state boundaries decrease the correlation coefficient without affecting the number of boundaries that align with a stimulus feature.

To be sure that we only correct for relevant covariates, a different set of covariates was used for each annotation of interest. Which covariates were chosen for which annotation of interest can be seen in Table 1. Thus, the residual of ‘small-scale location changes’ describes the moments at which there was a small-scale change in location, while correcting for low-level visual features, all 8 Alexnet layers, shots, MFCC, speech, and events. The various annotation categories can be divided into 5 groups: visual features (low-level visual features and Alexnet layers), locations (small-scale and large-scale), low-level audio (MFCC), mid-level audio (speech), and conceptual (event boundaries). We have chosen the covariates per annotation of interest such that the annotation is corrected for all annotations that are not in the same group, and not corrected for those within the same group, as such annotations share the same relevant information. For example, if we were to investigate the small-scale location changes, it would not make sense to correct for changes in large-scale locations as these annotations are nested.

**Table 1:** Overview of the covariates that were selected for each annotation category that was analyzed.

| Annotation of interest | Covariates |
| --- | --- |
| Small-scale locations | Low-level visual features |
| Large-scale locations | Alexnet layers |
|  | Shots |
|  | MFCC |
|  | Speech |
|  | Events |
| Low-level visual features | Small-scale locations |
|  | Large-scale locations |
|  | MFCC |
|  | Speech |
|  | Events |
| Speech | Low-level visual features |
|  | Alexnet layers |
|  | Shots |
|  | Small-scale locations |
|  | Large-scale locations |
|  | MFCC |
|  | Events |
| Events | Low-level visual features |
|  | Alexnet layers |
|  | Shots |
|  | Small-scale locations |
|  | Large-scale locations |
|  | MFCC |
|  | Speech |

The p-value that results from Pearson’s correlation between the residuals could not be used here as the degrees of freedom are incorrectly estimated for timeseries that are strongly auto-correlated by nature. We therefore used permutation testing to compute p-values. In each permutation, the order of neural states were shuffled within each run, while keeping the number of neural states and their duration intact. Then, partial correlation was again computed on the concatenated runs. In total there were 10,000 permutations to compute a null distribution per ROI and annotation category combination. The p-value was computed by counting how many values in the null distribution were above or equal to the actual measurement, and thus the test was one-tailed. All tests were corrected for multiple comparisons using Bonferroni correction, correcting for the number of ROIs.

As part of a more exploratory whole-brain analysis, we created 3461 partially overlapping spherical searchlights across the gray matter masked cortex. The centers of the searchlights were spaced 2 voxels apart, and the radius of each searchlight was set to 3.5 voxels. Only searchlights overlapping with at least 20 voxels were included, and the cerebellum and sub-cortical areas were excluded, giving us searchlights of up to 178 voxels (4806 *mm*3), with an average size of 144 voxels (3888 *mm*3). Then, the same analysis was applied to each searchlight as was done for the previously mentioned ROIs, except for the number of permutations, which was set to 1,000 per searchlight. To correct for multiple comparisons, we applied a false discovery rate (FDR) correction over all searchlights (Benjamini & Hochberg, 1995). A voxel was only interpreted as significant when the average p-value of all searchlights overlapping with that voxel was smaller than the highest p-value that survived the FDR-correction.

## 3 Results

### 3.1 Location changes are selectively associated with neural state boundaries in PPA and RSC

If neural states are driven by aspects in the environment to which a brain area is responsive, then the state boundaries in location-sensitive areas should align with moments in the movie at which there is a change in location. This association should remain present even when correcting for visual features at different levels of complexity, shots, speech, MFCC, and event boundaries. Indeed, the neural states boundaries in the right PPA showed a significant association with both small-(*p <* 0.001) and large-scale (*p <* 0.01) location changes (Figure 1A). For small-scale location changes we additionally found a significant alignment with neural states boundaries in the left RSC (*p <* 0.01), while the alignments in the left PPA (*p* = 0.010) and the right RSC (*p* = 0.019) did not survive the Bonferroni correction. The number of neural state boundaries was much higher in both PPA (right: 373, left: 367) and RSC (right: 412, left: 416) than the number of small-scale location changes (231 changes; see Figure S5), indicating that there is no one-to-one correspondence between neural states and locations. For large-scale location changes, the alignment with neural state boundaries in the left RSC (*p* = 0.019) did not survive Bonferroni correction. In absolute numbers, 24% of the changes in both small- and large-scale locations exactly aligned with a neural state boundaries in left PPA, and 26% of the changes in small-scale location and 28% of the changes in large-scale locations aligned with neural state boundaries in the right PPA. These numbers were similar in the RSC (left RSC: 26% for small-scale locations, and 27% for large-scale locations; right RSC: 24% for both small- and large-scale). See Figure S6 for a snippet of the changes in locations and neural state boundaries in PPA over time. The differences in results between the left and right ROIs could be related to the low number of subjects and consequently the low signal-to-noise ratio. Adding more subjects may alleviate the lateralization.

**Figure 1:**
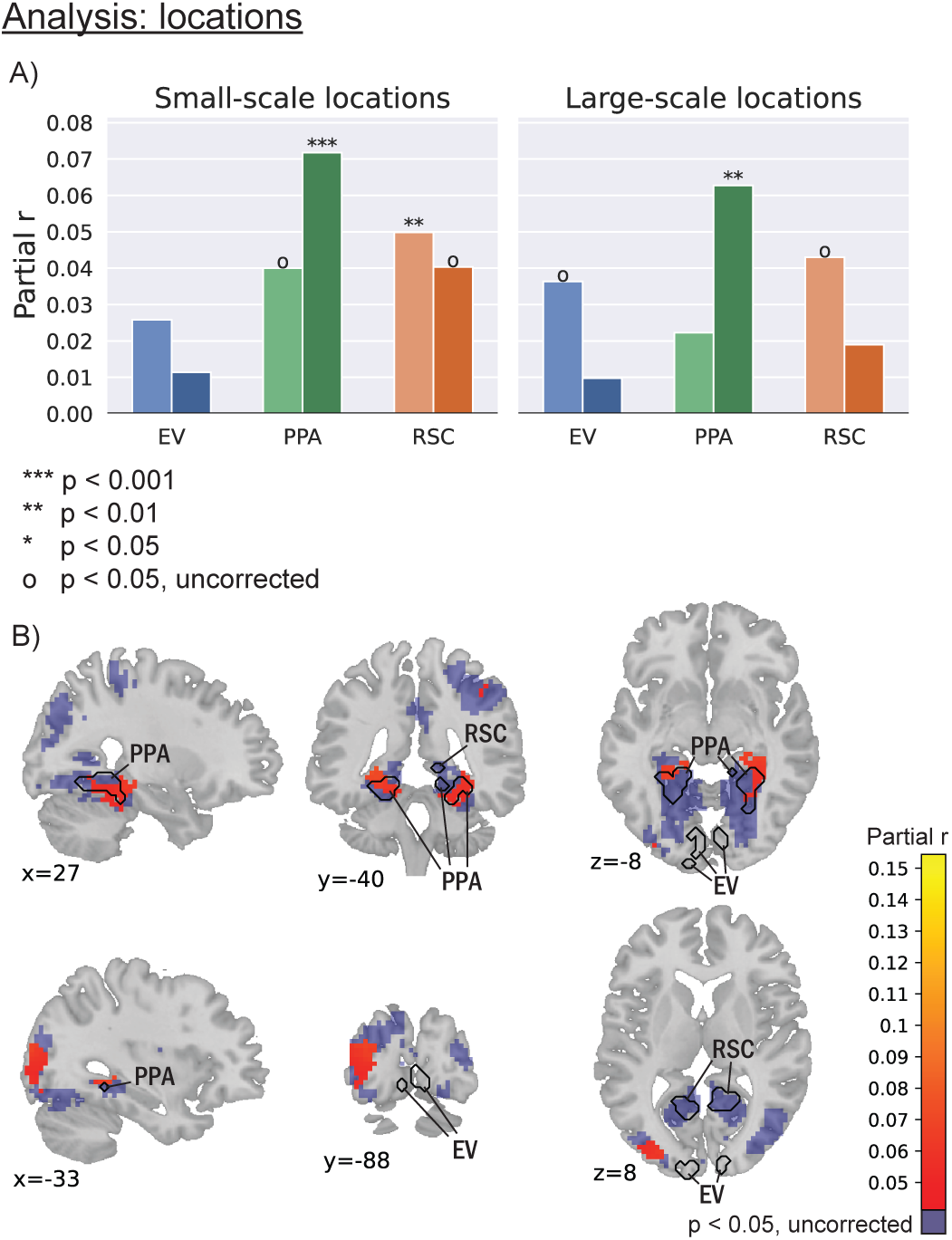
Neural state boundaries in the location-sensitive PPA and RSC showed a significant alignment with location changes, especially for the small-scale locations. Importantly, the alignment was absent in early visual areas. The values represent r values between neural state boundaries and changes in location, while correcting for low-level visual features, 8 layers of Alexnet, shots, MFCC, speech, and events. EV: early visual. PPA: parahippocampal place area. RSC: retrosplenial cortex. A) ROI analysis for both small-scale and large-scale locations. B) Thresholded results of the whole-brain analysis for small-scale locations. The results of the whole-brain analysis for large-scale locations can be found in Figure S8.

### 3.1 Location changes are selectively associated with neural state boundaries in PPA and RSC

Location changes are naturally accompanied by changes in visual features and also tend to occur when there is a new shot. To test whether our corrections for changes in low-level visual features were sufficient, we also investigated the left and right early visual areas, which should not show a significant alignment between their neural state boundaries and changes in location. In those ROIs, we indeed only observed a partial correlation with large-scale location changes in the left early visual (*p* = 0.027), which did not survive Bonferroni correction. These results suggest that the association we observed in the PPA and RSC cannot be explained by residual effects of changes in visual features, and that the association with locations is specific to the location-sensitive ROIs.

The specificity of the association between location changes and neural state boundaries was also supported by the exploratory whole-brain analysis. Two clusters with a significant alignment between the neural state boundaries and small-scale location changes were overlapping with both the left and right functionally defined PPA (Figure 1B, top). These clusters were also present for large-scale locations changes, though much smaller (Figure S8). A third small-scale location cluster was detected in left middle occipital gyrus (Figure 1B, bottom). This cluster may overlap with the occipital place area (OPA), which is an additional area that could be considered scene-sensitive (Dilks et al., 2013; Kamps et al., 2016). Indeed, when looking at the functional localizer results that were also used to define the PPA and RSC, we see an additional cluster where the OPA could anatomically be expected, and this cluster overlaps with our small-scale location cluster (Figure S7). Contrary to the ROI analysis, no location clusters overlapped with the functionally defined RSC. This may be because the correction for multiple comparisons was stricter in the whole-brain analysis than in the ROI analysis. Taken together, these results support the hypothesis that neural state boundaries overlap with location changes in location-sensitive regions specifically.

### 3.2 Changes in visual features align with neural state boundaries in visual areas

If changes in the relevant features in the environment are driving neural state boundaries, we should see this effect not only for location changes but also for other visual features. Therefore, we also looked at how changes in low-level visual features are associated with neural state boundaries in the early visual areas, while correcting for locations changes, MFCC, speech, and event boundaries. Indeed, we observed that both the left and right early visual areas showed a significant alignment between neural state boundaries and changes in low-level visual features (*p <* 0.001; see Figure 2A). This alignment was also observed in left and right PPA (*p <* 0.01), but the correlations in left RSC did not survive Bonferroni correction (*p* = 0.020). The clusters found in the exploratory whole-brain analysis suggested a more regionally specific association between changes in visual features and neural state boundaries. A cluster was present in medial V1 (Figure 2B, top), overlapping with the functionally defined early visual areas. A second cluster was present in the left frontal eye fields (FEF; Figure 2B, bottom). Contrary to the ROI analysis, no cluster overlapped with the functionally defined PPA or RSC, which may be because the correction for multiple comparisons was stricter in the whole-brain analysis than in the ROI analysis. We did indeed observe alignment between neural state boundaries and changes in low-level visual features in the PPA (but not the RSC) in the whole-brain analysis when no correction for multiple comparisons was performed. Another possibility is that the difference in shape is detrimental: both left and right functionally defined PPA and RSC had an elongated shape (Figure S3), while the searchlights used in the whole-brain analysis are round and defined by a center and a radius, possibly making the whole-brain analysis unable to capture the full characteristics of the PPA and RSC.

**Figure 2:**
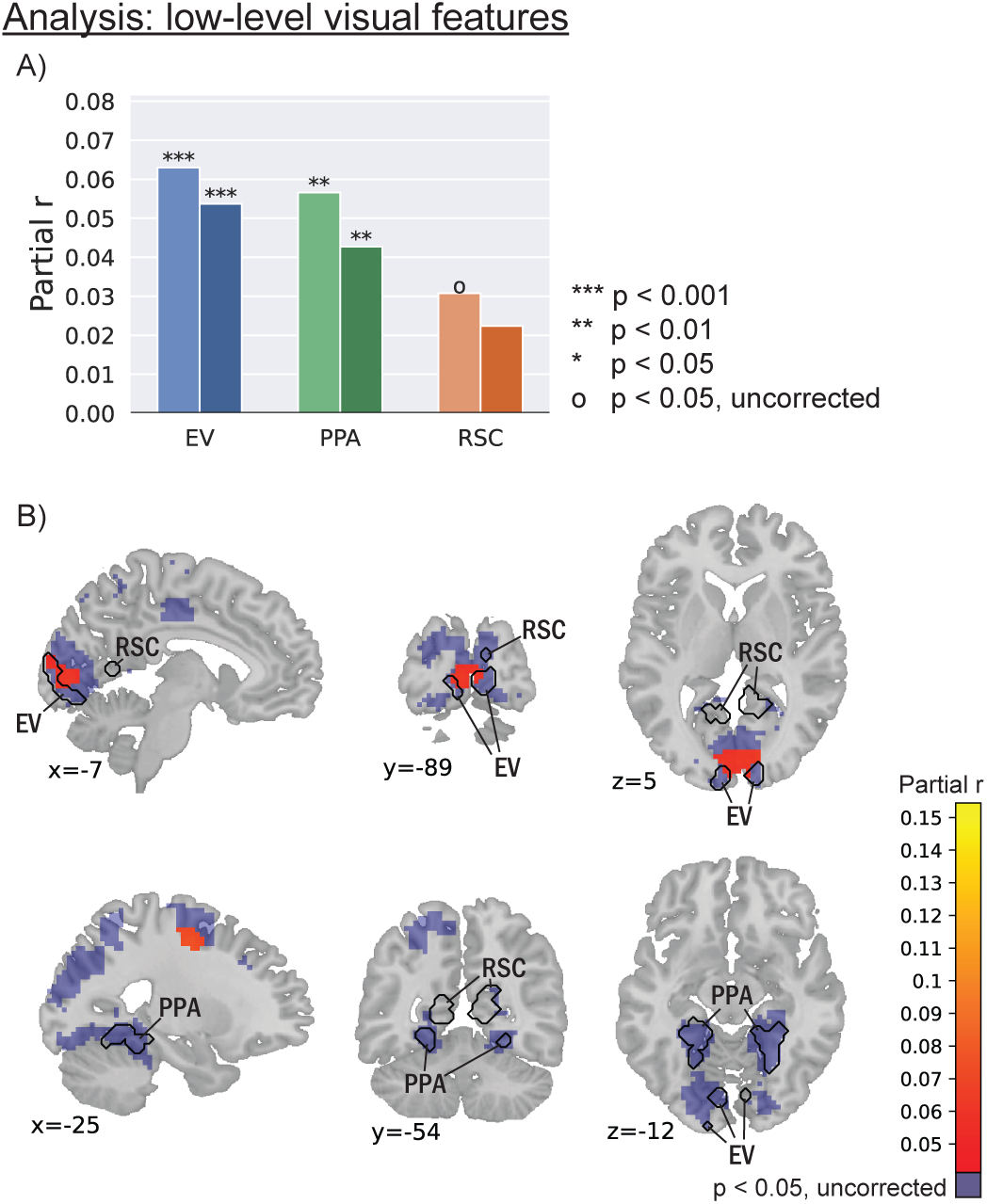
Neural states in early visual areas showed a significant alignment with low-level visual features, in both the ROI and the whole-brain analysis. In the ROI analysis, this alignment was also present in the PPA. The values represent r values between neural state boundaries and changes in low-level visual features, while correcting for small- and large-scale locations, MFCC, speech, and events. EV: early visual. PPA: parahippocampal place area. RSC: retrosplenial cortex. A) ROI analysis. B) Thresholded results of the whole-brain analysis.

### 3.3 Alignment between neural state boundaries and the external environment is also present in the auditory domain

To further investigate the generalizability of neural states aligning to specific features in the stimulus, we additionally tested the alignment with speech onsets and offsets across the whole brain to study the auditory domain, while correcting for low-level visual features, Alexnet layers, shots, small- and large-scale locations, MFCC, and events. Many of the voxels with a significant alignment were centered around the bilateral temporal cortex (Figure 3), and were specifically found in bilateral middle temporal gyrus, superior temporal gyrus, temporal pole, Heschl’s gyrus, insular cortex, central and parietal operculum cortex, planum temporale, supplementary motor cortex, supramarginal gyrus and pre- and postcentral gyrus. Many of these areas are related to language comprehension (see e.g., Friederici & Gierhan, 2013). Moreover, additional regions were specifically found in the left hemisphere and included inferior frontal gyrus (i.e., Broca’s area), middle and superior frontal gyri, and the angular gyrus.

**Figure 3:**
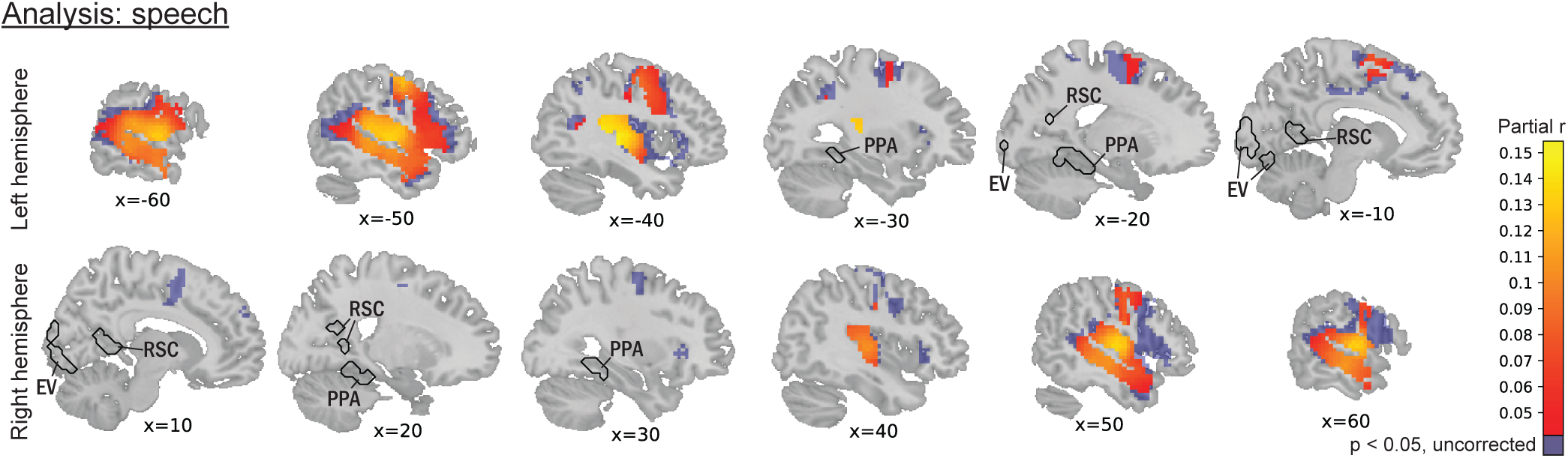
Voxels with a significant alignment between neural state boundaries and on- and offset of speech were found in various language-related regions, including bilateral middle temporal gyrus, temporal pole, Heschl’s gyrus, and left inferior frontal gyrus. The values represent r values between neural state boundaries and on- and offsets of speech across the brain, while correcting for low-level visual features, 8 layers of Alexnet, shots, small- and large-scale locations, MFCC, and events. EV: early visual. PPA: parahippocampal place area. RSC: retrosplenial cortex.

### 3.4 Event boundaries align with neural states in higher-level cortical areas

Alignments between neural state boundaries and event boundaries have been investigated before in previous literature, but not yet while correcting for visual and auditory covariates in the stimulus. To investigate to which extent these alignments could be driven by visual or auditory aspects rather than the conceptual features related to event segmentation, we performed our whole-brain analysis on event boundaries as well, while correcting for visual features at various levels of complexity, as well as for shots, MFCC, speech, and location changes. The clusters found to show a significant alignment with event boundaries were numerous and spread throughout the cortex (Figure 4), and could be found in some high-level areas such as the angular gyrus, precuneus, inferior frontal gyrus, prefrontal cortex, and posterior cingulate cortex, but not in low-level visual areas or primary auditory cortex.

**Figure 4:**
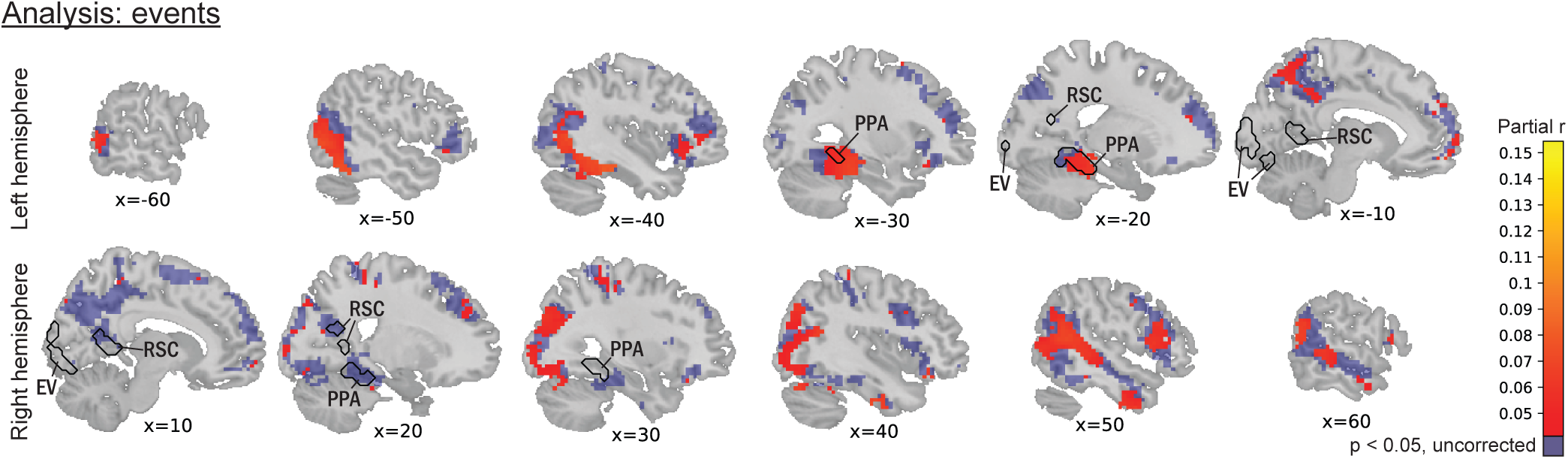
Voxels with a significant alignment between neural state boundaries and event boundaries were found throughout the brain, including the angular gyrus and prefrontal cortex, but excluding low-level visual areas and primary auditory cortex. Values represent r values between event boundaries and neural state boundaries across the brain, while correcting for low-level visual features, 8 layers of Alexnet, shots, small- and large-scale locations, MFCC, and speech. EV: early visual. PPA: parahippocampal place area. RSC: retrosplenial cortex.

### 3.5 Correcting for covariates is crucial to isolate feature specific associations

Because of the high correlations in the various annotation categories, we used partial correlation in our analysis to minimize the possibility that a significant correlation could be explained by a covariate rather than the annotation of interest. To investigate the consequences of this approach, we also performed the same analysis in Supplementary Section F but with Pearson’s correlation rather than partial correlation. In summary, we found that many areas now showed a significant alignment with more annotation categories than in the partial correlation analysis, in both the ROI analysis and the whole-brain analysis. In particular for events, the vast majority of the cortex now showed a significant alignment. This suggests that the correction for covariates allowed us to better isolate feature specific associations between neural state boundaries and changes in the stimulus.

## 4 Discussion

Understanding how the brain segments information as it receives continuous and dynamically varying sensory input is of critical importance for moving to a more ecologically valid understanding of brain function (Lee et al., 2020; Willems et al., 2020). Recent work has established that event segmentation could be underpinned by neural states, which form a temporal hierarchy across the cortex. The aim of this study was to test whether these neural states reflect meaningful temporal units in the environment. Our findings provide support for this hypothesis and suggest that the cortical temporal hierarchy found in previous literature partly reflects the temporal scales at which representations in the environment evolve. In the following paragraphs we will first discuss the evidence for this conclusion in more detail before looking at broader implications of this research for our understanding of neural state segmentation in the brain.

### 4.1 Neural states at different levels of the cortical hierarchy are shaped by aspects of the environment to which these brain areas are responsive

It has previously been hypothesized that neural states should reflect meaningful units in the environment (Baldassano et al., 2017). However up to now, neural state boundaries have only been linked to event boundaries specifically (Baldassano et al., 2017; Geerligs et al., 2022; Mariola et al., 2022) and it has not been investigated if neural states at different levels of the temporal hierarchy indeed represent different meaningful units. In line with this hypothesis, we found that there is a significant alignment between changes in location and neural state boundaries in the place-sensitive PPA and RSC. The whole brain analysis showed that this alignment was highly specific to such place-sensitive areas. Importantly, these significant alignments could not be explained by changes in visual features of various levels of complexity, nor by MFFC or speech, nor by event boundaries. Additional support was found by investigating neural states in early visual areas, which did not show this association with locations, but did have a significant alignment with low-level visual features. Again, the whole brain analysis revealed that this association between low-level features and neural states was highly specific to early visual areas. Besides this alignment in the visual domain, we also found some support in the auditory domain: neural state boundaries in various language-related areas showed an alignment with the onset and offsets of speech, which cannot be explained by MFCC and various other covariates. Together these results provide strong support for our hypothesis that neural states in a particular brain area reflect the stable representations of a feature that is relevant to that area, and present in the external environment.

Although we found alignments between the ends of neural states and the ends of meaningful units in the stimulus, neural states and stable features in the environment do not have a one-to-one correspondence. In general, brain regions are not necessarily uniquely responsive to one specific measurable aspect in the environment (Fedorenko et al., 2013; Woolgar et al., 2016). In our study in particular, there were many more neural states boundaries in both the PPA and the RSC than the number of location changes throughout the stimulus, indicating that one neural state does not correspond to one location on the screen. In the PPA specifically, a significant association between neural states and low-level visual features was found in addition to the association with locations. Previous studies have found that the PPA is sensitive to high spatial frequencies (Rajimehr et al., 2011) in addition to places (Seeliger et al., 2021), and texture (Cant & Goodale, 2011), while some even argue that the PPA could consist of multiple subregions, with one subregion specialized in low-level visual features and another in scenes (Baldassano et al., 2013), or with one biased to static stimuli and another to dynamic stimuli (Çukur et al., 2016). Thus, neural states within a particular brain area could pertain to multiple aspects of the environment, making it difficult to relate a specific neural state to one specific feature in the stimulus.

### 4.2 Neural states are more than passive representations

In this study, we focused on the idea that neural states are a reflection of the environment. However, they are likely to be more than just a passive representation of the current sensory input. Instead they could also be associated with how we act on that environment, what we know about the environment from previous experience, and what we are attending to (Baldassano et al., 2017; Zacks & Tversky, 2001). For example, our exploratory whole-brain analysis revealed a cluster in the FEF that had a significant alignment between neural states and low-level visual features. An increase in FEF activity has previously been related to saccades around event boundaries (Rosano et al., 2002; Speer et al., 2003). Others have more recently found that gaze patterns change after event boundaries (Eisenberg & Zacks, 2016), as well as showing an increase in similarity across participants (Smith et al., 2024). These saccades partially happen because event boundaries tend to co-occur with changes in low-level visual features. Thus, our cluster in the FEF probably demonstrates an association between neural state boundaries and eye movements, indicating that active processes, such as eye movements, could also be associated with neural states.

Evidence for the idea that neural states might reflect what we know about the environment from previous experience comes from previous literature showing that neural states may contain information that is currently not accessible in the environment, while still being relevant. For example, Zadbood et al. (2017) showed that the same event-specific activity pattern occurs while experiencing a stimulus, while remembering that same stimulus, and while listening to another person describing it. Thus, activity patterns can be specific to a stimulus even in its absence. Additionally, Ezzyat and Davachi (2021) found object-specific activity to be persistent even after the object had changed, but only as long as the event had not ended yet. This persistent activity pattern seemed to reflect a stable mental model within the event. Thus, neural states could reflect the current model or representation of an event or context (DuBrow et al., 2017; Richmond & Zacks, 2017).

Some brain areas, such as early sensory areas, may be more sensitive to the information that is currently present in the environment, while others may rely more on previous input (Hasson et al., 2008). This possibility is partially reflected in our results, where the low-level visual features only reflect information that was available on the screen, with a significant alignment in early visual areas. In contrast, event boundaries are much more conceptual and require knowledge about the situation beyond the current sensory information, and these event boundaries aligned with neural state boundaries in higher-level areas. By having neural states across the cortex, it is possible to have many such representations with different levels of abstraction across many different time scales. These multi-scale representations could aid information processing by reducing complexity at each scale, and together representing a multi-scale environment (Quax et al., 2020). Given the high number of areas that have been reported to be related to events or event boundaries, different parts of the brain may be representing different aspects of event information.

In line with the notion of neural states, previous literature suggests that there might be a ‘gating’ mechanism that separates distinct mental representations (Chien & Honey, 2020). Information from the recent past is either integrated or not, making it possible to separate currently relevant information from interfering information (Ezzyat & Davachi, 2011; Kurby & Zacks, 2008; Shin & DuBrow, 2020; Zacks et al., 2007). Here we showed that the timing of neural state boundaries is meaningful in multiple brain areas, providing additional support that the gate closes at appropriate moments. Which moments are considered to be appropriate, could be influenced by top-down mechanisms such as attention, as neural states are affected by the task at hand (De Soares et al., 2024).

### 4.3 The temporal cortical hierarchy is partly driven by the temporal evolution of various features in the environment

Neural state durations have been shown to form a nested temporal hierarchy (Baldassano et al., 2017; Geerligs et al., 2022), in line with previously found TRWs (Hasson et al., 2008). Our finding that these neural states align with meaningful units in the environment, suggests that the neural state hierarchy could naturally arise from how fast various aspects of information from the environment tend to evolve: low-level visual features can change very quickly, while a change in location occurs much less often. Previously reported variations in the duration of neural states could be explained by a variation in duration of the relevant aspect in the external environment, such as a movie scene at one location taking longer than a scene at another location (Geerligs et al., 2022). Moreover, a nested hierarchy of information is additionally present in the environment and could be reflected in nested neural states: when there is a change in location, there will always be a change in low-level visual features, but not vice versa. Taken together, we show that the hierarchical information that is present in the natural environment is at least one possible component that contributes to the temporal cortical hierarchies that have been reported in previous literature. However, it is likely that this is just one of multiple aspects that contribute to the hierarchies, as intrinsic timescale and power spectra gradients have been reported in the absence of a stimulus as well (Honey et al., 2012; Raut et al., 2020; Stephens et al., 2013).

In the context of event cognition, previous theories have proposed different ways in which event representations could be updated. Some theories propose that event representations are updated incrementally (e.g., Event Indexing Model (Zwaan et al., 1995); Huff et al. (2014)). This would entail that a change in spatial location without a change in characters only affects the part of the event representation that relates to the location, and not the part that relates to the characters. Alternatively, event representations could be updated globally, meaning that a change in location affects the event representation as a whole, including non-changing characters (e.g., Event Segmentation Theory; Zacks et al., 2007). Event representations could also be updated both incrementally and globally (e.g., Gernsbacher’s Structure Building Framework (Gernsbacher, 1997); Bailey and Zacks (2015), Kurby and Zacks (2012)), suggesting that some changes result in incremental updates, while others bring about global updates. Based on our findings, we can now say that we see evidence for incremental local updating in the brain, given the specific areas of significant alignment between neural state boundaries and changes in location. An overall global neural updating across the cortex is unlikely, given that the alignment with event boundaries (after correction for some covariates) is not found across the whole cortex, but rather only in clusters. However, conclusions about cognitive global event representations are beyond the scope of the current study as neural states and event perception are two separate but possibly related concepts. Instead, we can now start building a bridge between neural states and descriptives of event cognition and behavior.

### 4.4 The merits and drawbacks of naturalistic stimuli

Naturalistic stimuli such as movies and audio books contain a lot of complex information, with different features simultaneously developing over time at multiple scales. Together, these evolving features form a coherent narrative. This contributes to their ecologically validity, as the same type of information is present in the real-world environment (Willems et al., 2020). Studying the brain in an ecologically valid environment facilitates the possibility to study the brain during an engaging and realistic experience (Lee et al., 2020), and is essential to prevent oversimplified theories (Cantlon, 2020; Nastase et al., 2020).

Multi-scale information that is present in both the real-world environment and naturalistic stimuli often forms a hierarchy, with for example a new person entering the room coinciding with a major change in visual and/or auditory information. Therefore, when annotating multiple aspects in a naturalistic stimulus, such aspects are likely to be highly correlated (Hamilton & Huth, 2020; Nastase et al., 2020). To deal with this issue, we corrected for a multitude of covariates in our analyses, including visual features (to study location-specific states), auditory features, and event boundaries. We observed much more regionally specific associations between neural states and different annotation categories when a correction for covariates was performed, compared to when no corrections were applied. This can be illustrated by looking at the association between event boundaries and neural states. Some of the regions that showed associations between neural states and events in previous work (Geerligs et al., 2022), including early visual regions, anterior insula, and anterior cingulate cortex, only showed a significant alignment between neural state boundaries and event boundaries in the absence of any correction for covariates. This suggests that this association may be because of the co-occurrence between event boundaries and visual or auditory changes, rather than essentially being associated with events. However, most higher-level regions that were observed before, also showed an association between neural states and events with corrections for covariates. These regions included part of the cingulate cortex, part of the prefrontal cortex, part of the middle frontal gyrus, precuneus, middle temporal gyrus, and superior temporal gyrus, suggesting that these areas contain representations of abstract and gist-level descriptions of an event (Bird, 2020).

Even though our correction for covarying visual features was effective, it may not have been perfect. For example, when looking at the association between low-level visual features and neural states in the whole-brain analysis, we observed an initially unexpected cluster in left FEF. Instead of concluding that the FEF is sensitive to these low-level visual features, it would be more reasonable to relate this alignment to eye movements. These results highlight that even though we carefully selected a broad set of covariates, we can never be completely confident that we were able to account for all covarying features. Indeed, since for example location changes can only be observed by combining many different visual features in the environment, a full disentanglement may be impossible. Although this sounds like a drawback of naturalistic stimuli, it is also a reflection of their ecological validity, as those correlations are inherently present in the real world as well. This issue becomes even more complicated when investigating an abstract concept such as event boundaries. We did find various clusters around the brain that show an alignment with event boundaries, but we cannot conclude that these exact areas are involved in (the processing of) event representations. Instead, some of these areas may align with a currently unknown stimulus feature, or even internal representations or thoughts that simply correlate with event boundaries. Furthermore, such event boundaries reflect a change in one or multiple dimensions of the stimulus, while some of these dimensions (e.g., a change in character, object, action, or goal; Zacks et al., 2009) can also be considered part of what it means to have an event boundary. If we would account for all possible dimensions as covariates, including high-level information such as a change in the goal of a character, no information of the event would be left. Therefore, the covariation of various aspects of the same stimulus is an important aspect to keep in mind when analyzing naturalistic data, and including a correction for covariates in the analyses is one way to disentangle some of these effects. The covariates should then be chosen carefully: if one would like to claim that a certain observation (e.g., neural correlates to changes in location) cannot be explained by a covariate (e.g., low-level visual features), this should be corrected for. However, especially with more abstract information (e.g., event boundaries), a correction for a covariate that diminishes the definition of such information (e.g., a change in a character’s goal or action) should not be included.

### 4.5 Conclusion

In conclusion, we found support for the hypothesis that neural state boundaries in a specific area occur when an aspect in the environment that is relevant to that area changes. This also suggests that neural states reflect stable features in the environment and provides further support for the idea that neural state segmentation at different levels of the cortical hierarchy underlies event segmentation, aiding information processing as well as memory. Finally, our results indicate that the cortical temporal hierarchy in terms of neural state durations partly reflects the temporal scales at which different aspects of the environment naturally evolve.

## Data and Code Availability

In this study we made use of open data from different sources.

*•* General description of the StudyForrest dataset: studyforrest.org.
*•* Manually denoised movie fMRI data: openneuro.org/datasets/ds001769/versions/1.2.2.
*•* Anatomical MRIs: github.com/psychoinformatics-de/studyforrest-data-structural.
*•* Functional localizer fMRI data: github.com/psychoinformatics-de/studyforrest-data-aligned.
*•* Stimulus annotations of shots and locations: github.com/psychoinformatics-de/studyforrest-data-annotations.
*•* Stimulus annotations of low-level visual features: github.com/psychoinformatics-de/studyforrest-data-confoundsannotation.

All code for further data preprocessing and analysis can be found here: github.com/dynac-lab/stable-features-reflected-in-neural-states

## Ethics

This study utilized open data from StudyForrest. According to the accompanying literature (Hanke et al., 2016; Hanke et al., 2014; Sengupta et al., 2016), all participants were fully instructed about the nature of the study, gave their informed consent for participation as well as for public sharing of all obtained data in anonymized form, and received monetary compensation. Ethical approval was obtained from the Ethics Committee of the Otto-von-Guericke University

## Author Contributions

Djamari Oetringer: Conceptualization, Methodology, Software, Formal analysis, Visualization, Writing - Original Draft. Dora Gözükara: Methodology, Software, Writing - Review & Editing. Umut Güçlü: Conceptualization, Resources, Writing - Review & Editing, Supervision. Linda Geerligs: Conceptualization, Methodology, Resources, Writing - Review & Editing, Funding acquisition, Supervision.

## Acknowledgements

We would like to thank Aya Ben-Yakov for providing us with the event segmentation results of the StudyForrest movie stimulus.

## Supplementary Material

Supplementary Material (created during production as a web link to online material).

