## Supplementaries for "The Neural Basis of Event Segmentation: Stable Features in the Environment are Reflected by Neural States"

### Supplementary Material

#### A Annotation descriptives

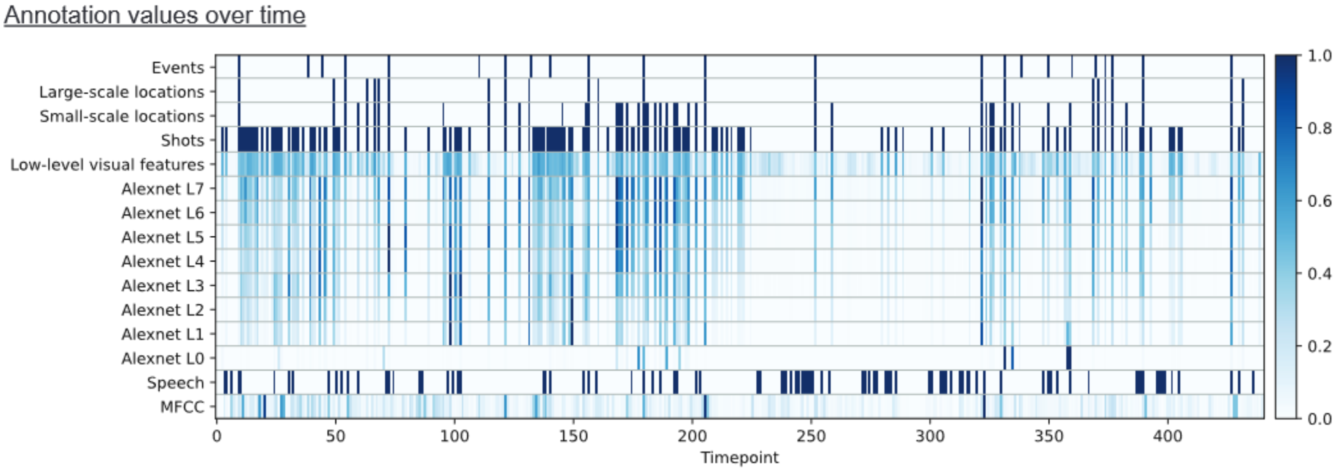


**Fig. S1.** Annotation timeline of run number 2, including a delay of 4.5 seconds to account for the haemodynamic response delay, and after downsampling.

Figure S1 visualizes all annotations over time of one particular run. Notably, many but not all event boundaries align with location changes and shots. The second and third event boundaries in this segment (timepoints 38 and 44) do not seem to align with substantial changes in sensory inputs. Instead, they correspond to pivotal moments in the narrative. Namely, the main character is running away from bullies while his legs are supported by leg braces. At the second event boundary of the run, these braces start to fall apart. At the third event boundary of the run, the main character starts running independently without any braces.

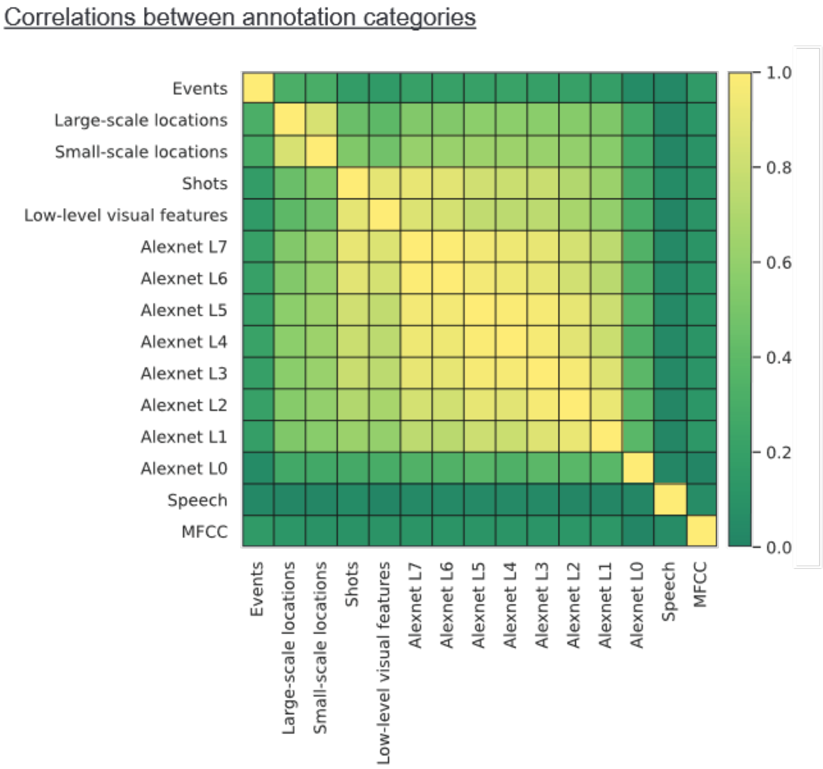


**Fig. S2.** Pearson's correlation between all the annotation categories after downsampling to fMRI volumes. The number after "Alexnet L" indicates the layer of Alexnet, where "L0" is the input layer.

Figure S2 visualizes the various correlations between all annotation correlations used in this study. Overall, speech had the lowest correlation with an average correlation coefficient of 0.10, followed by MFCC with 0.17. Despite event boundaries being likely to occur with changes in location, the correlation between these annotations were only 0.30 (small-scale) and 0.32 (large-scale). This may partially be because event boundaries were identified based on button presses, thus some time passes before participants realize that a new event has started and press the button. How much time is necessary can vary between event boundaries, as some transitions are more apparent than others. Therefore, some event boundaries co-occur with for example changes in location, while others seem to be slightly delayed.

#### B ROI descriptives

**Table S1.** Descriptives of the functionally defined ROIs.

| ROI | Threshold (t) | MNI center (x y z) | | | Number of voxels | Volume ($mm^3$) |
|---|---|---|---|---|---|---|
| Early visual left | 3.50 | -10.44, | -94.26 | 0.84 | 100 | 2700 |
| Early visual right | 3.39 | 9.96 | -90.33 | 3.45 | 100 | 2700 |
| PPA left | 4.50 | -22.68 | -44.31 | -11.64 | 100 | 2700 |
| PPA right | 5.79 | 25.17 | -44.22 | -12.60 | 100 | 2700 |
| RSC left | 4.36 | -13.60 | -57.78 | 10.21 | 77 | 2079 |
| RSC right | 4.55 | 14.04 | -51.78 | 9.06 | 100 | 2700 |

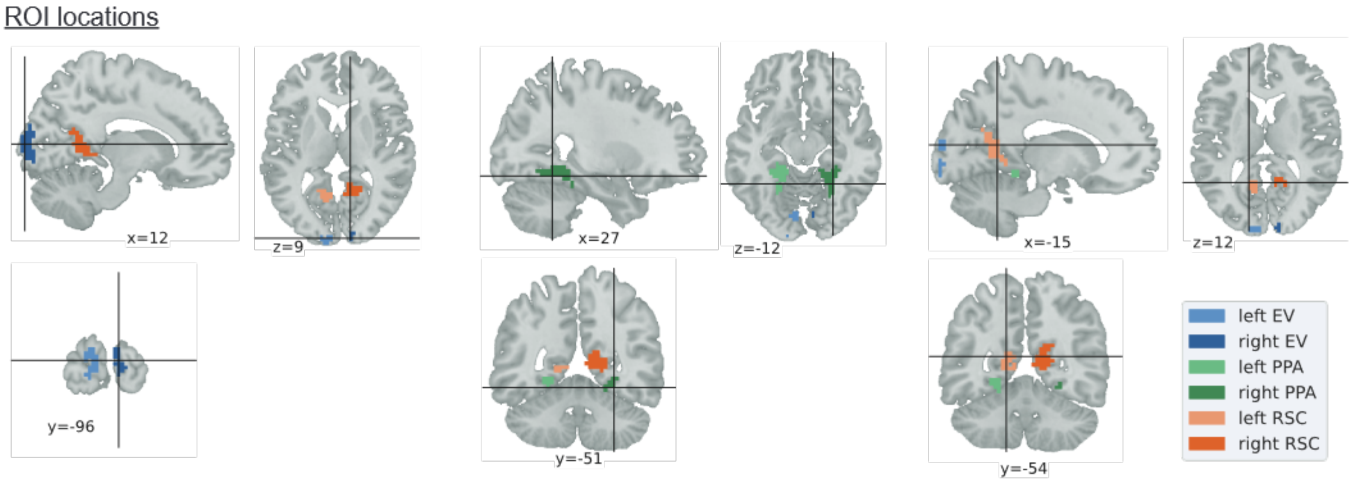


**Fig. S3.** Location of each functionally defined ROI. EV: early visual. PPA: parahippocampal place area. RSC: retrosplenial cortex.

Chosen thresholds, MNI coordinates, and sizes of all ROIs can be found in Table S1. The shape and location of each ROI is visualized in Figure S3. The left RSC only had 77 voxels rather than 100, as a slightly higher t-value gave a region of 193 voxels. This is because there were two clusters connected to each other with just one voxel.

#### C Hemodynamic response delay

To compute the appropriate hemodynamic delay for aligning neural state boundaries and movie annotations, we assumed that the neural state boundaries in all ROIs are correlated with event boundaries in the stimulus, as previously found by Baldassano et al. (2017) and Geerligs et al. (2022). For each ROI, we extracted the neural states boundaries of run 4, which was not used in any other analysis, and set all timepoints with a boundary to 1 and all others to 0. We then computed the Jaccard index between these neural state boundaries and the event boundaries for a given delay. The delay was varied between 0 and 10 s with steps of 0.5 s, which was applied before downsampling the event boundaries from seconds to fMRI volumes. This analysis is in line with Geerligs et al. (2022), who showed that the optimal delays across the cortex were similar to previously found BOLD signal delays. The results of our analysis can be found in Figure S4. Based on these results, we chose to use a delay of 4.5 s, as this was the optimal delay for the left and right PPA and the right early visual, in between two optimal delays of the left RSC, and close to the optimal delay of the right RSC. Left early visual however is an outlier and had an optimal delay of 6 s.

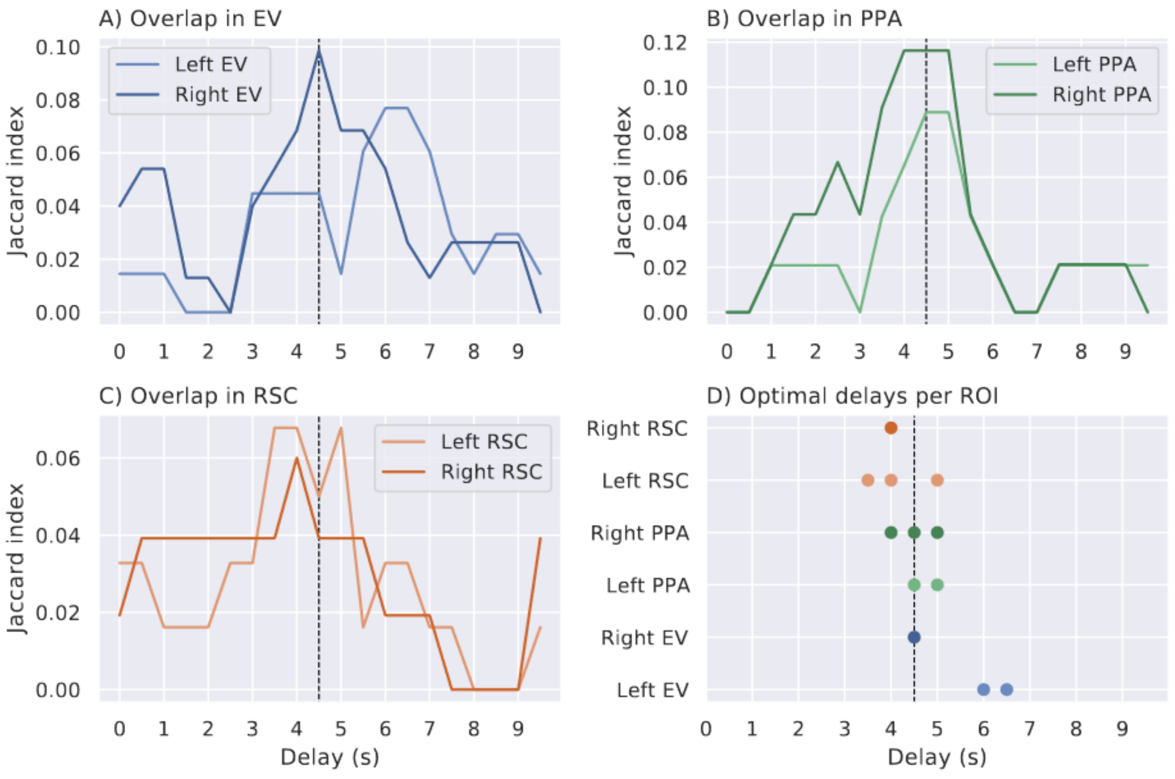


**Fig. S4.** Jaccard index between the event boundaries and the neural state boundaries of run 4 over various delays. A,B,C) the Jaccard index over delay for each ROI. D) the optimal delay per ROI. The dashed lines at 4.5 s show the delay that we used in the main analysis. EV: early visual. PPA: parahippocampal place area. RSC: retrosplenial cortex.

#### D Number of neural state boundaries and annotation boundaries

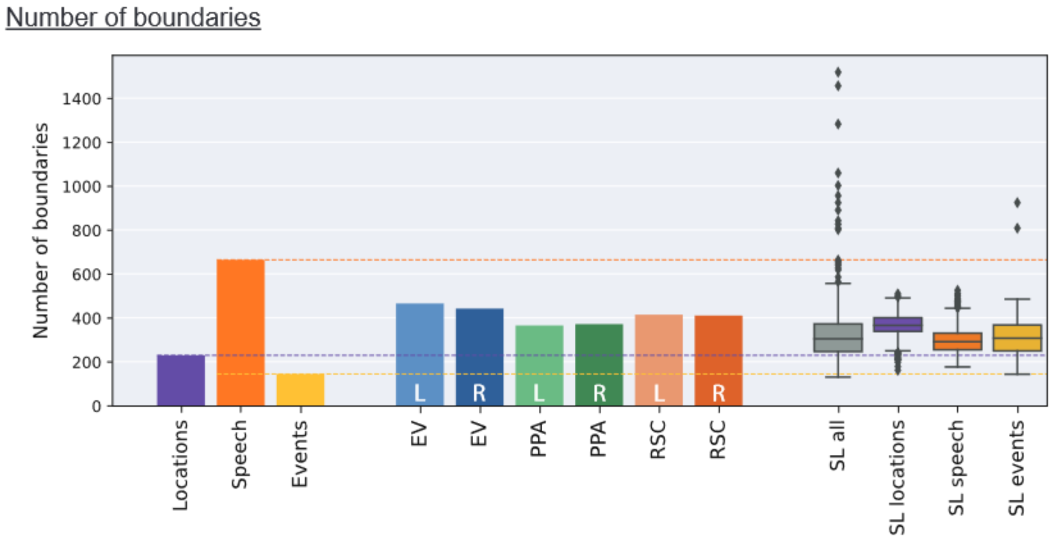


**Fig. S5.** Throughout the movie stimulus, 231 changes in small-scale locations, 666 speech onsets/offsets, and 146 event boundaries occurred (excluding run number 4). All ROIs and many searchlights had a higher number of neural state boundaries as compared to the number of changes in location and event boundaries, and less than the number of speech onsets/offsets. Locations: small-scale locations. EV: early visual. PPA: parahippocampal place area. RSC: retrosplenial cortex. SL all: all searchlights across the brain. SL locations/speech/events: searchlights with a significant alignment between their neural states and small-scale locations/speech/events, after FDR-correction.

Figure S5 summarizes the number of small-scale location changes, speech onsets/offsets, event boundaries, and neural state boundaries across the movie stimulus, excluding run number 4. In total, there were 231 changes in small-scale location, 666 onsets/offsets of speech, and 146 event boundaries. The number of changes in location and the number of event boundaries are much lower than the number of neural state boundaries in each ROI (left early visual: 467, right early visual: 444, left PPA: 367, right PPA: 373, left RSC: 416, right RSC: 412). When taking all searchlights throughout the cortex, the number of neural state boundaries per searchlight ranged from 132 to 1520. When only selecting those searchlights that had a significant alignment between locations and neural states, 270 out of 279 searchlights had a higher number of neural state boundaries than the number of changes in location (231), with a minimum of 163 and a maximum of 511. For searchlights with a significant alignment with speech, all searchlights had fewer neural state boundaries than the number of onsets/offsets in speech, ranging from 178 to 527. When only selecting searchlights with a significant alignment with events, only one searchlight had fewer neural state boundaries (145) than number of event boundaries (146), and the remaining searchlights ranged from 152 to 926 neural state boundaries. Although many areas showed an alignment between their neural states and locations or events, there were often more neural state boundaries than moments in the movie stimulus with a relevant change. This indicates that neural states in specific areas reflect more than just a stable unit of one particular feature.

#### E Additional descriptives of location analysis

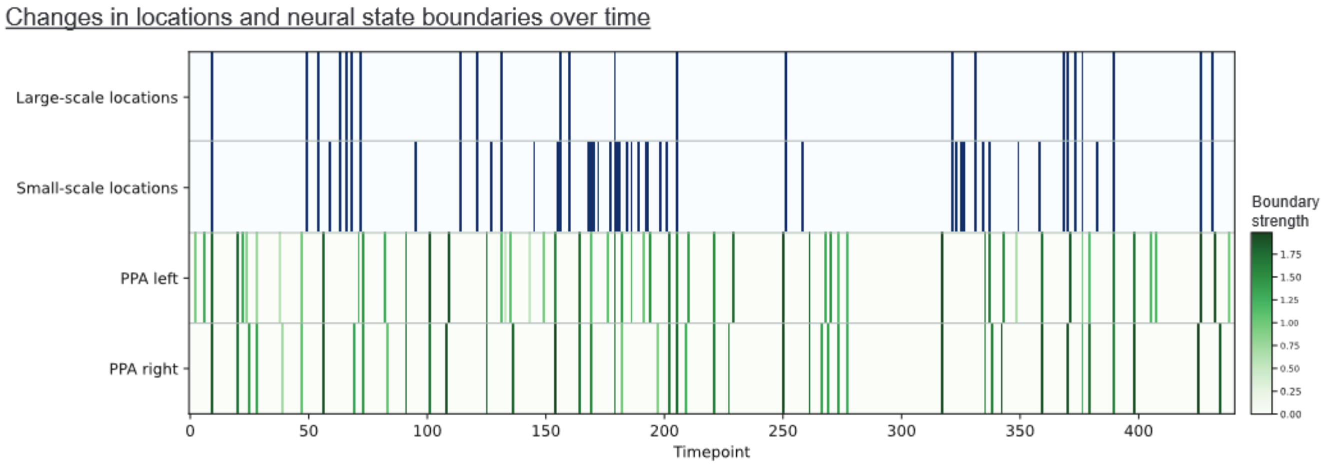


**Fig. S6.** Large-scale and small-scale location changes have a nested pattern. When visually inspecting these annotations and the neural state boundaries in the PPA, some overlap seems to be present but there are many neural state boundaries without an accompanying change in location. PPA: parahippocampal place area.

Figure S6 visualizes the changes in location over time for run number 2, together with the neural state boundaries in left and right PPA. The boundaries between left and right PPA look similar. Additionally, some overlap between location changes and neural state boundaries is visible, but there are many neural state boundaries without a change in location.

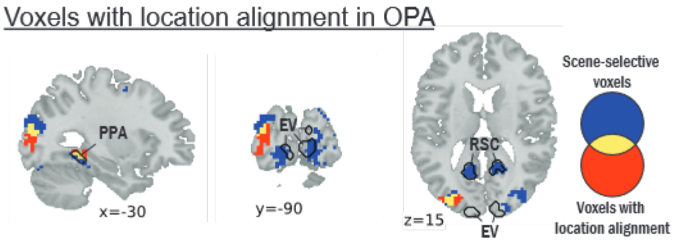


**Fig. S7.** Voxels that show a significant alignment between changes in small-scale locations and their neural state boundaries overlap with thresholded contrast of houses and landscapes with all other image categories (used to define PPA and RSC) of the functional localizer. The threshold is set to $p = 0.001$ ($t = 3.79$). EV: early visual. PPA: parahippocampal place area. RSC: retrosplenial cortex.

The whole-brain results of small-scale locations (Figure 1) revealed an initially unexpected cluster in the occipital cortex, which could actually be overlapping with the OPA. To test this, we took the results of the between-subject t-test per voxel that was also used for the definitions of PPA and RSC. As can be seen in Figure S7, when thresholding at $p = 0.001$, the cluster showing an alignment with locations indeed overlaps with a scene-sensitive cluster that could be considered the OPA.

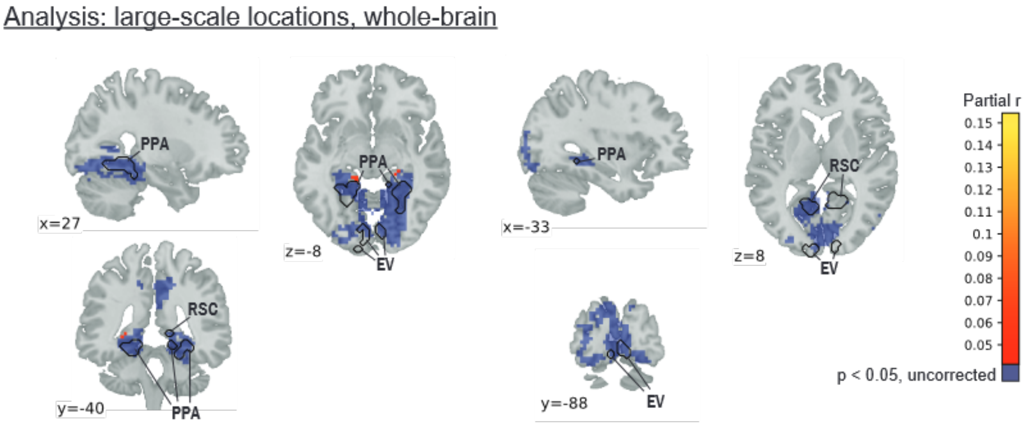


**Fig. S8.** Voxels with a significant alignment between neural state boundaries and large-scale location changes were sparse and could be found close to the PPA, but these clusters were small. Values represent r values between neural state boundaries and large-scale locations across the brain, while correcting for low-level visual features, 8 layers of Alexnet, shots, MFCC, speech, and events. EV: early visual. PPA: parahippocampal place area. RSC: retrosplenial cortex.

In the main text we showed the association between neural states boundaries across the brain and small-scale location changes (see Figure 1B). Here, we also visualized the association with large-scale location changes (Figure S8). For large-scale locations, some voxels with a significant alignment could be found close to the PPA. Compared to the whole-brain analysis with the small-scale locations, the clusters overlapping with the PPA were much smaller, and the cluster in left OPA disappeared.

#### F Results without correction for covariates

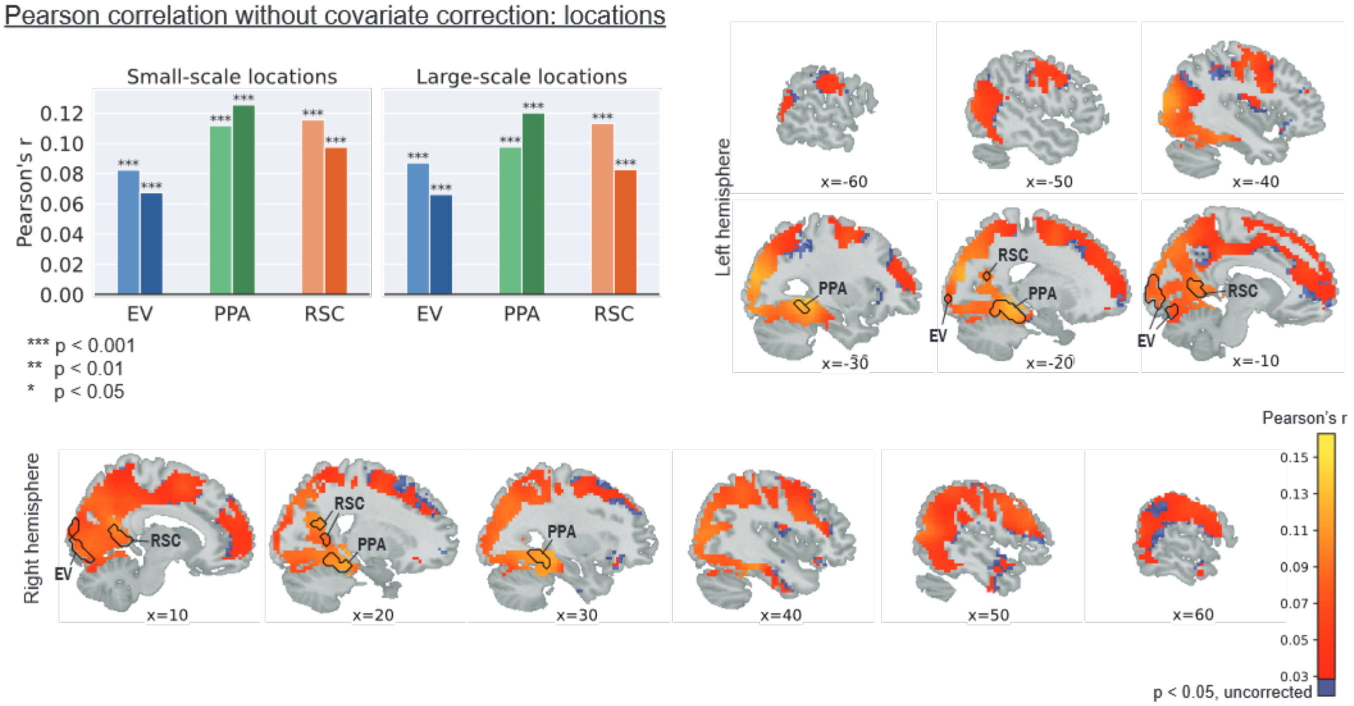


**Fig. S9.** When not correcting for covariates, neural states in all ROIs and in many areas across the cortex showed a significant alignment with locations. The values represent Pearson's r between neural state boundaries and changes in location. EV: early visual. PPA: parahippocampal place area. RSC: retrosplenial cortex. The brain plots only show the results of the whole-brain analysis for small-scale locations. The results of the whole-brain analysis without covariate correction for large-scale locations are not depicted here.

To investigate the importance of correction for covariates, we also reran our main analyses with Pearson's correlation as the measurement rather than partial correlation. For changes in location, neural state boundaries now showed a significant alignment in all ROIs ($p < 0.001$), and this alignment was present across the vast majority of the cortex, including early visual areas as well as prefrontal areas (Figure S9). A similar pattern was found in the alignment between neural state boundaries and changes in low-level visual features (Figure S10): this alignment was significant in all ROIs ($p < 0.001$). Furthermore, voxels in which neural state boundaries aligned with changes in low-level visual features now extended across higher-level areas, with some clusters being found in the frontal areas and the middle cingulate cortex. The clusters of significant alignment between event boundaries and neural state boundaries were larger compared to the analysis with covariates, with significant voxels now also being present in low-level visual areas (Figure S12). In contrast, the voxels with a significant alignment between neural states and speech without any correction for covariates (Figure S11) were very similar to those voxels with a significant alignment after correcting for a set of covariates (Figure 3): only 688 voxels were added as compared to a differences of more than 10,000 voxels for the other annotation categories.

Taking all annotation categories together, without any correction for covariates many areas showed overlap between two or more categories, with many visual areas showing a significant correlation between neural state boundaries and changes in low-level visual features, location changes, and event boundaries. Overall, regions that showed significant alignment in our partial correlation analysis still had a significant alignment when not correcting for any covariates, but many additional regions now showed the same alignment. Thus, correcting for covariates is necessary to exclude the possibility that the significant alignment found in a specific region or voxel is actually related to another aspect of the movie.

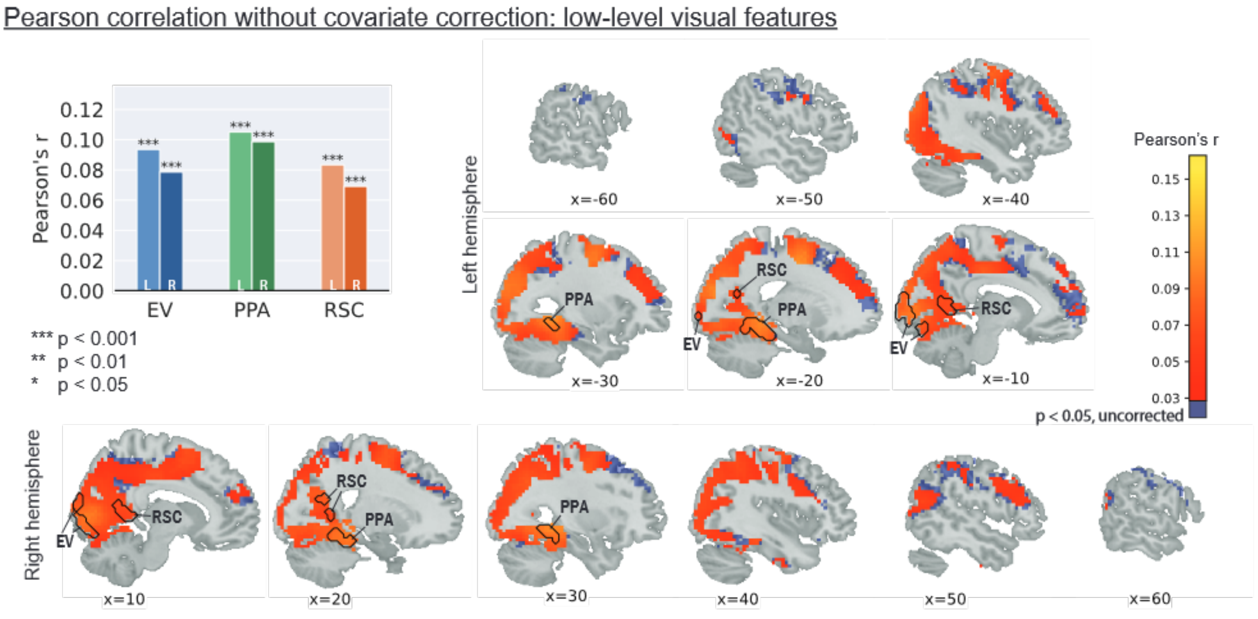


**Fig. S10.** When not correcting for covariates, neural states in all ROIs and in many areas across the cortex showed a significant alignment with low-level visual features. The values represent Pearson's r between neural state boundaries and changes in low-level visual features. EV: early visual. PPA: parahippocampal place area. RSC: retrosplenial cortex.

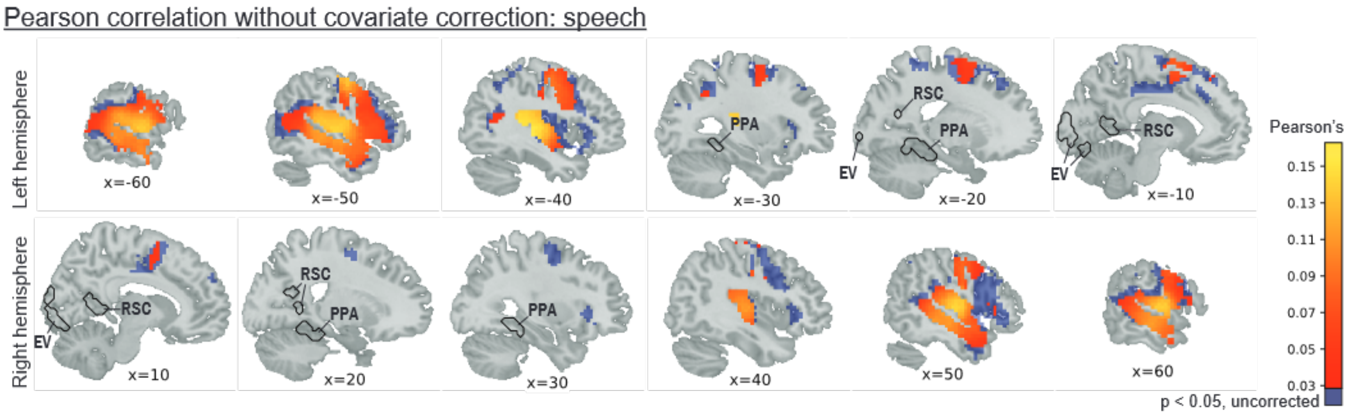


**Fig. S11.** When not correcting for covariates, voxels with a significant alignment between speech onsets and offsets are relatively similar to those voxels that are found when correcting for a set of covariates (Figure 3). The values represent Pearson's r between neural state boundaries and speech. EV: early visual. PPA: parahippocampal place area. RSC: retrosplenial cortex.

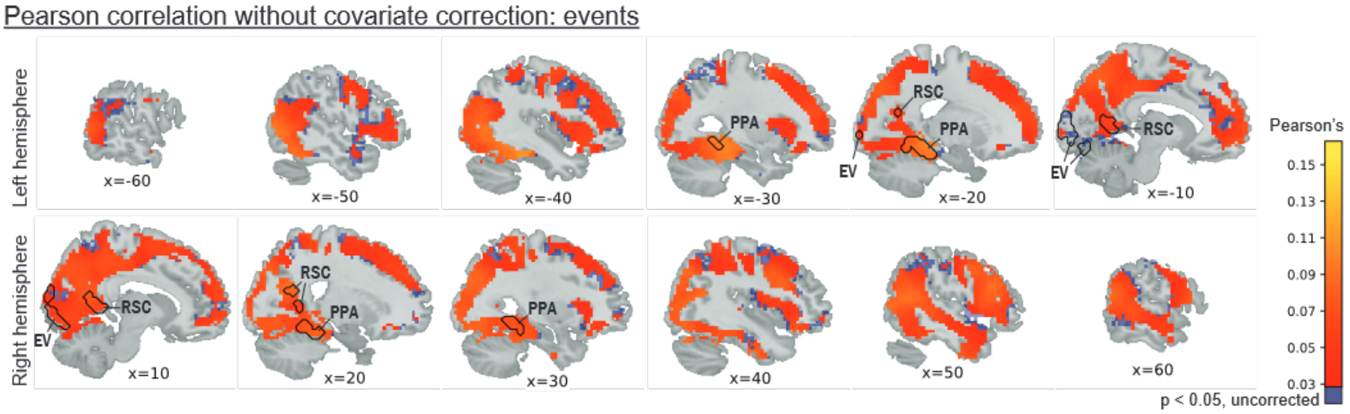


**Fig. S12.** When not correcting for covariates, neural states in all ROIs and in many areas across the cortex showed a significant alignment with events. The values represent Pearson's r between neural state boundaries and event boundaries. EV: early visual. PPA: parahippocampal place area. RSC: retrosplenial cortex.
